# Interactions between immunotoxicants and parasite stress: implications for host health

**DOI:** 10.1101/194142

**Authors:** Ross D. Booton, Ryo Yamaguchi, James A. R. Marshall, Dylan Z. Childs, Yoh Iwasa

## Abstract

Many organisms face a wide variety of biotic and abiotic stressors which reduce individual survival, interacting synergistically to further reduce fitness. Here we studied the effects of two such synergistically interacting stressors; immunotoxicant exposure and parasite infection. We model the dynamics of a within-host infection and the associated immune response of an individual. We consider both the indirect sub-lethal effects on immunosuppression and the direct effects on health and mortality of individuals exposed to toxicants. We demonstrate that sub-lethal exposure to toxicants can promote infection through the suppression of the immune system. This happens through the depletion of the immune response which causes rapid proliferation in parasite load. In addition, high toxicant exposure can alter cellular regulation and cause the breakdown of normal healthy tissue, from which we infer higher mortality risk of the host. We classify this breakdown into three phases of increasing toxicant stress, and demonstrate the range of conditions under which toxicant exposure causes failure at the within-host level. These phases are determined by the relationship between the immunity status, overall cellular health and the level of toxicant exposure. We discuss the implications of our model in the context of individual honey bee health. Our model provides an assessment of how pesticide stress and infection interact to cause the synergistic breakdown of the within-host dynamics of individual honey bees.

## 1. Introduction

During their lifetime, organisms are exposed to a wide range of chemical, physical and biological stressors. Exposure to environmental (e.g. anthropogenic, climatic) and natural stress (e.g. pathogens, parasites and predation) reduces individual fitness [1]. Recently, there has been increasing interest in multiple stress approaches, examining the potential for stressors to interact synergistically, defined as the combined effects of stress having a greater impact than expected [2]. Understanding the mechanisms behind these synergistic interactions is important for quantifying the true impacts of individual anthropogenic stress on organisms [3].

Pesticides are an important class of anthropogenic toxicant stress, with the use of pesticides continuing to increase globally [4, 5, 6]. Pesticides are seen as crucially important to crop productivity, preserving around one-fifth of total crop yield contributing to food security [7]. Concerns about the detrimental impacts of these pesticides [8, 9] have in the past forced policy makers to restrict the application of some insecticides [10]. Non-target insects frequently encounter these insecticides [5], with concentrations able to build up throughout food sources and within various life-stages of the organism [11, 12, 13, 14, 15, 16, 17].

Toxicants such as pesticides can cause lethality [18, 19, 20], but more often have other sub-lethal effects such as impairments on foraging [21, 22, 23, 24], feeding [25], learning [26, 27], memory [28, 27] and fecundity [29, 30, 31]. Exposure during early life can have both lethal and sub-lethal effects later appearing during adulthood [32, 33]. These environmental contaminants can interact synergistically in combination with other natural stressors. For example, combinations of toxicant exposure with parasite infections can increase individual mortality [34, 35], increase the initial pathogen load [36, 37] and increase virulence [38]. Synergistic toxicant-pathogen interactions have been observed in many types of organisms such as insects, snails, water fleas, frogs, salamanders, fish and mussels (see review by Holmstrup et al., 2010). In addition to toxicants causing direct lethality, they can also cause indirect damage to individual immune defence. Individual organisms defend them-selves against various infections via a suite of immune responses, and these can be damaged or inhibited through toxicant exposure [39]. For example, pesticides have been shown to reduce the total hemocyte abundance in insects [40, 41], the nodulation initiation [40, 42], the encapsulation response [43, 41] and antiviral defences [44].

Of particular recent concern are the widespread losses to global wild and managed honey bee populations [6, 45, 46]. The Western honey bee (*Apis mellifera* L.) is widely recognised as the most important commercial insect pollinator [47, 48, 49, 50], contributing to global food security and biodiversity [51, 52]. While a single cause for these widespread colony losses has yet to be identified, there is agreement that it may have its origins within multiple stressors interacting with each other [53, 54, 55, 56]. Possible candidates include neonicotinoid pesticides [12, 13, 57, 27], mites [58, 59], viruses [60, 61, 62] and microsporidia infections [63, 64].

In this study, we examine the general mechanism by which immunotoxicants interact with infection to reduce host health. This observed synergy between multiple stressors is currently poorly understood from an immunological perspective [65]. We focus our study on the general ecotoxicological applications of the model, in the case of any immunotoxicant interacting with any parasite infection. We do this by formulating a system of nonlinear ordinary differential equations (ODEs) to investigate the consequences of immunosuppression by a toxicant and the effect this has on within-host infection. We first consider a toxicant-free environment to examine the conditions under which the infection can spread. We then consider the interaction between the infection and both lethal and sub-lethal exposure to toxicants and examine the outcome on within-host dynamics. We also consider the case of aggressive direct lethality of toxicants on the production of new tissue cells.

## 2. The Model

The immune response of any individual relies upon the interdependent defence of physical, humoral and cellular responses, denoted in our model by immune function *Z*. Nowak and May [66] proposed a general model to describe the interaction between a cellular immune response and a replicating virus, in the setting of self-regulating cytotoxic T lymphocytes (CTLs) targeting infected cells. The model they present is simple but captures the fundamental biological processes governing the immune response to foreign antigens, and following this framework we denote within-host cell density as *X*. We denote the total parasite/pathogen density as *Y*. The total number of cells within the model represents a general susceptible subset of animal tissue cells. As a motivating example, our model can be thought of describing the midgut epithelial cells of the honey bee *X* under a *Nosema ceranae* infection *Y* [67] with associated immune response *Z*, although we also propose that our model can be thought of describing any interaction between any immunotoxicant and associated parasite or pathogen in a general host.

Toxicants can be lethally toxic to individuals at high enough exposure [18, 19, 20]. In addition various functions associated with the immune response are damaged by toxicants [39, 40, 41, 42, 43, 68, 69, 70, 44]. We model both the direct lethality (denoted by parameter *r*) and indirect sub-lethal immunotoxicity (denoted by parameter *h*) effects of toxicant exposure *Q*. For simplicity, we assume fast dynamics of virus replication compared to the replication of other immune or within-host cells resulting in the formulation of the model (Figure 1) as a 3-compartmental set of nonlinear ODEs;

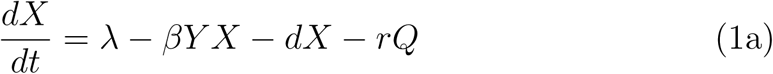

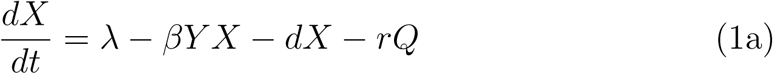

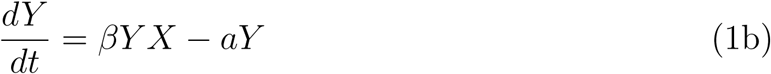

with *c - hQ >* 0 and *λ – rQ >* 0. When *Z* = 0 (the immune response is depleted), we remove equation (1c) from system (1) and the system becomes the two dimensional system of equations (1a) and (1b) without the immune response term –*pY Z*;

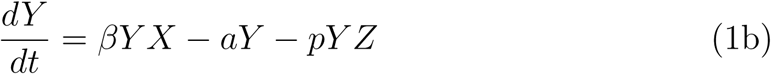

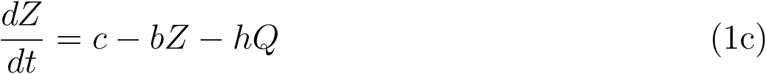

**Figure 1:**
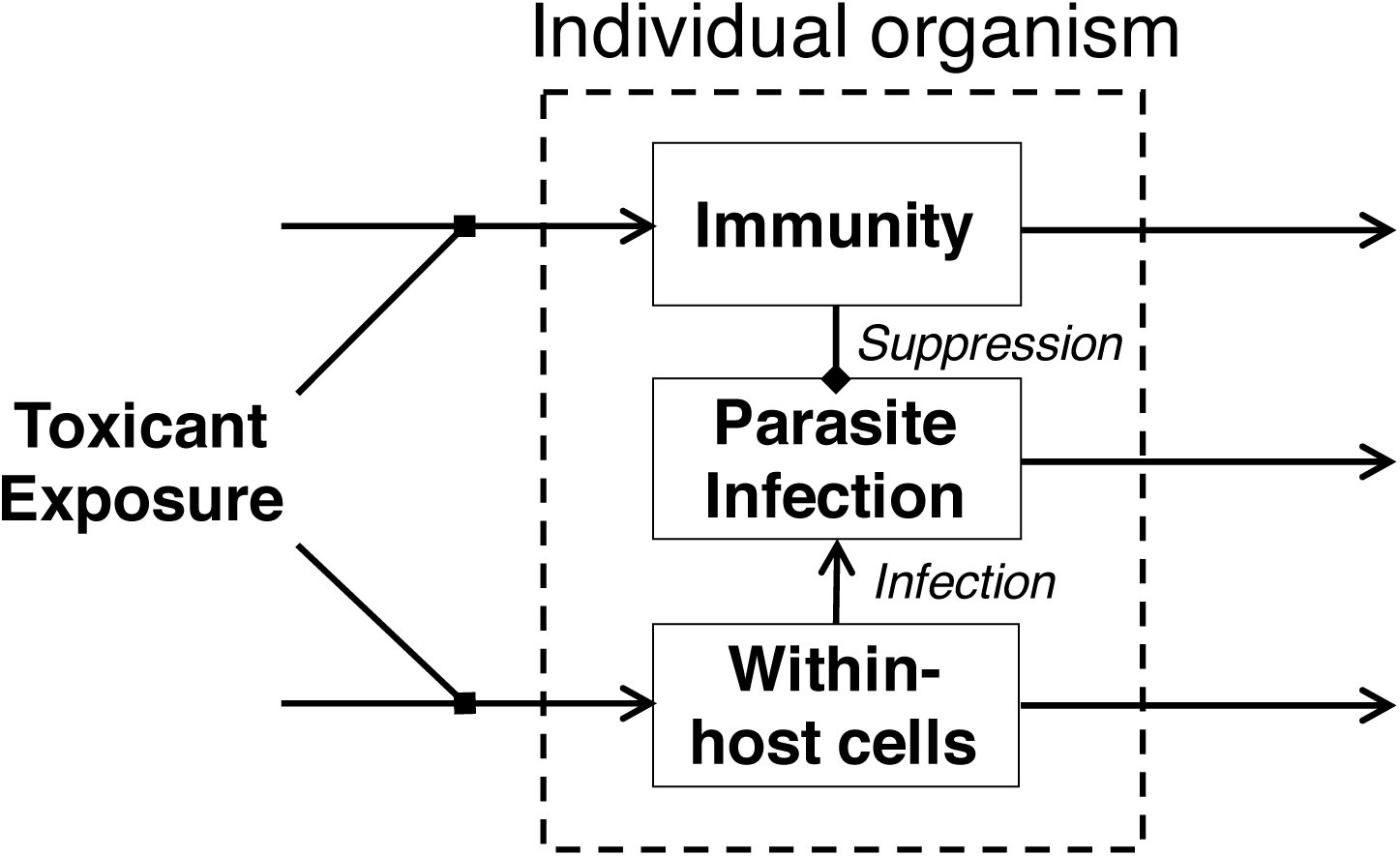
The modelling framework we use to model the interaction between toxicant exposure and parasite infection in an individual. Block arrows represent suppression. We model toxicant exposure as a suppressive effect on immunity and within-host cells

We assume that within-host cells are produced at rate *λ*, and die at percapita rate *d*. Parasites are created at rate *β* via a linear mass action, and are removed at per-capita rate *a*. The immune response *Z* is activated upon encountering parasites *Y* and the removal of parasites occurs at rate *α*. Although in reality, functions involved in immunity are not activated on the instance of meeting the parasite, but there is a complicated intermediary chain between processes which eventually result in the removal of parasites [71]. For simplicity, we assume that this complicated process can be summarised by our function *pY Z*. We assume that the immune dynamics *Z* are decoupled from those of within-host and parasite density. This represents the simplest possible assumption and various extensions to this assumption are possible. Immunity is therefore produced at rate *c*, and is removed at per-capita rate *b*.

Within our model we infer the mortality risk of the host through the status of the within-host cells *X*, so that individual mortality risk is high when the number of cells *X* is small. This condition enables us to think about the mortality risk of an individual analogous to a highly infected within-host tissue (e.g. parasite infection within the gut of a honey bee).

Our system of equations (1) were analysed using standard stability methods from dynamical systems theory and solved numerically with Wolfram Mathematica version number *10.0.2.0*, using parameters taken from Table 2. We performed a full parameter dependence analysis which demonstrated the same two universal behaviours of the model which enabled us to choose arbitrary parameter sets.

**Table 1:** The net change of immunity, within-host cells and parasites after the introduction of toxicant, compared to the no-toxicant model, for both initial and no parasite infection load.

|  | No parasite infection | Initial parasite infection |
| --- | --- | --- |
| Within-host cells $X$ | reduced by $\frac{rQ}{d}$ | reduced by $\frac{hpQ}{b\beta}$ |
| Parasites $Y$ | no change | increased by $\frac{bQ(hp\lambda - abr - cpr)}{(ab+cp)(ab+p(c-hQ))}$ |
| Immunity $Z$ | reduced by $\frac{rQ}{d}$ | reduced by $\frac{hQ}{b}$ |

**Table 2:** The parameters used in the analysis of the model.

| Parameter | Symbol | Value |
| --- | --- | --- |
| production of within-host cells | $\lambda$ | 0.1 |
| rate of parasite infection | $\beta$ | 0.01 |
| death of within-host cells | $d$ | 0.01 |
| direct lethal effect of toxicant | $r$ | 0.1 |
| toxicant exposure | $Q$ | $[0, 1.5]$ |
| death rate of parasites | $a$ | 0.01 |
| immune suppression | $p$ | 0.009 |
| production of immunity | $c$ | 0.1 |
| removal of immunity | $b$ | 0.02 |
| indirect sub-lethal effect of toxicant | $h$ | 0.3 |

## 3. Results

In the following section we consider the baseline case of parasite infection in a toxicant-free environment before analysing our within-host system under the addition of a toxicant. We then consider the absence of direct lethal effects of toxicants before presenting the unique case of an aggressive toxicant.

### 3.1. Toxicant-free model

Initially we examine system (1) under the condition of the absence of toxicant exposure (denoted by subscript A). Two possible outcomes are possible. First the infection is removed entirely by the immune system, in which case the total within-host cells and total immunity each reach a constant level at the disease free equilibrium (DFE):

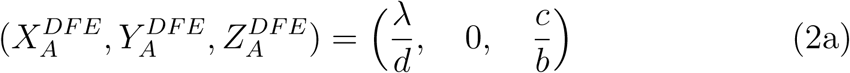

Where 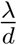 and 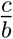 represent the ratio of total production to total removal of both within-host cells and immunity in the absence of toxicant respectively. Secondly the model predicts that an individual can become infected with parasites (*Y >* 0) under the following endemic equilibrium (EE):

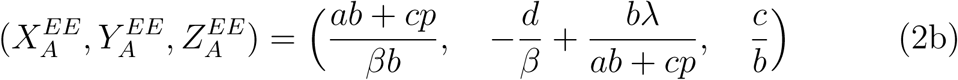

This shows that it is possible for an individual honey bee to sustain a partial parasite infection without the addition of any toxicant in our model. The expression 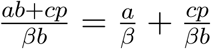 represents the reduction in within-host cells.

### 3.2. Toxicant-Parasite model

Next we consider system (1) under the condition of an infection and toxicant exposure (denoted by subscript B). In this case the model predicts two possible outcomes. First, the parasite infection is removed either by immune suppression or by the direct effects of the toxicant on the production of within-host cells represented by the DFE:

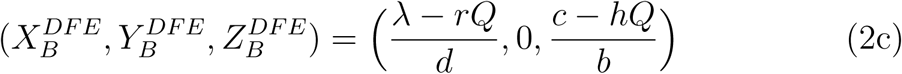

so that the addition of any toxicant reduces the total within-host cells by 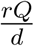 and reduces the immune function by 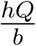. Secondly the model predicts an infected individual under toxicant exposure represented by the EE:

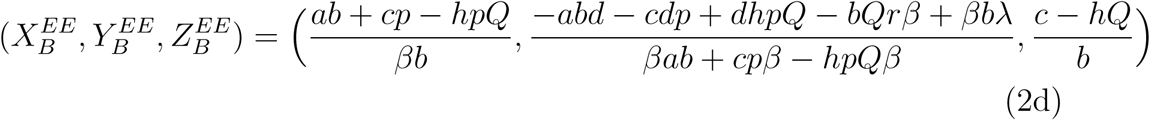

In this case, the parasite density grows rapidly as a result of the toxicant suppressing the immune system. The introduction of the toxicant reduces both within-host cells and immunity in both an infection-free and infected individual, but an initial parasite infection is required for an infection to grow.

The effect of toxicant exposure on the net change of within-host cells, parasite density and immunity within the individual is summarised in Table 1.

Next we assume that the indirect (sub-lethal) effects of toxicant exposureon immunosuppression are more prominent than the direct (lethal) depletion of within-host cells. With an initial infection *Y >* 0 we define this as occurring when the immune status of an individual is destroyed before the infection is removed or when

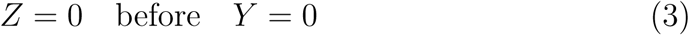

We summarise the behaviour of the model under this condition (Figure 2) into 3 distinct phases which describe the mechanism underlying the interaction between toxicant exposure and infection at the within-host level of the organism, and the parameter dependence of infection and immunity at equilibrium. Note that the total number of cells within an individual organism is not constant. This is because both parasite and within-host cells are removed by either the toxicant exposure or infection and new cells are produced.

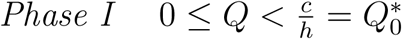

**Figure 2:**
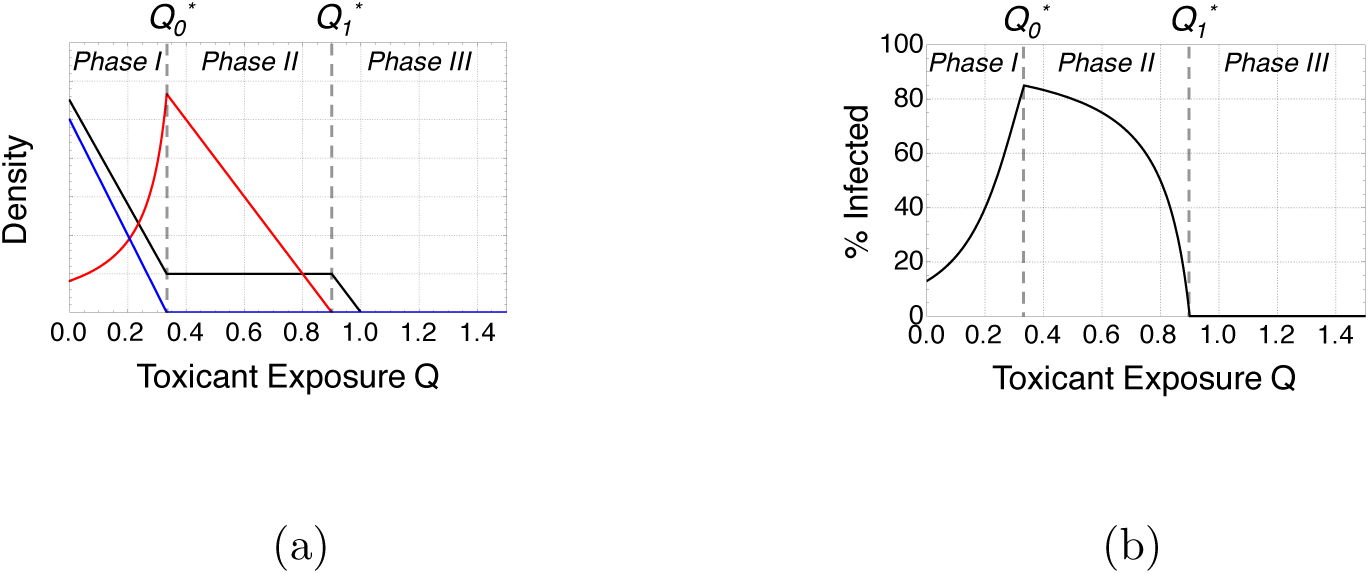
The mechanism of parasite infection under increasing toxicant exposure. This shows the parameter dependence of immunity, parasite density and within-host cells at equilibrium within the dynamics of our model. In (a) the total densities of immune function (blue), parasite load (red) and within-host cells (black) change as an individual is subject to higher toxicant loads. In (b) the total % parasite infection (black) changes as the toxicant load is increased. Parameters as in Table 2.

The model predicts that the initial state of an immune response is able to counter any infection. However, as the toxicant load is increased, the immune system is gradually depleted. Through a weakened immune suppression, this enables the parasite density to increase.

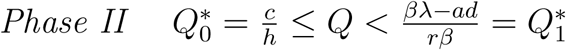

The second phase begins at the point of maximum infection and where the immune system has been completely inhibited. The increase in toxicant stress gradually depletes the parasite density while the within-host cells remain constant.

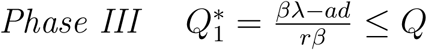

In phase three, the immune system has been destroyed and the parasite infection is no longer present leaving only a small fraction of within-host cells. Finally, the lethality of the toxicant causes the mortality of the individual honey bee and production of new cells ceases.

Thus we have calculated the conditions under which the within-host dynamics change according to the level of toxicant exposure. By understandingthe relationship between the parameters in the model and toxicant stress, we can make some biological interpretations. We predict that the ratio of the production of immunity to the amount of immunotoxicity 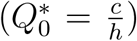 determines the point at which the infection load is at a maximum. The expression can be thought of as an indicator of immune status, and the point at which the toxicant stress becomes equal 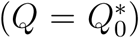 represents the complete inhibition of the immune system. The expression 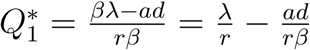 represents the point at which the ratio of cell production to lethal toxicant mortality (indicator of within-host cell status) compares to the ratio of the loss of cells to the toxicant cell depletion multiplied by the transmission of the infection. Therefore this condition represents the status of within-host dynamics and can be thought of as an indicator of health. When 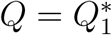 the infection has been removed but the overall health status is very low, from which we infer a higher mortality risk of the host. Therefore we have conditions describing how toxicant exposure relates to that of the immune status 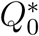 and overall health 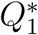 of the organism.

Our model predicts that a small amount of toxicant can cause the outbreak of an otherwise controlled infection. A healthy immune response can suppress the parasite infection to a very low level (Figure 3.a), but a small amount of toxicant can cause the status of both infection-free and infected individuals to decline rapidly (Figure 3.b).

**Figure 3:**
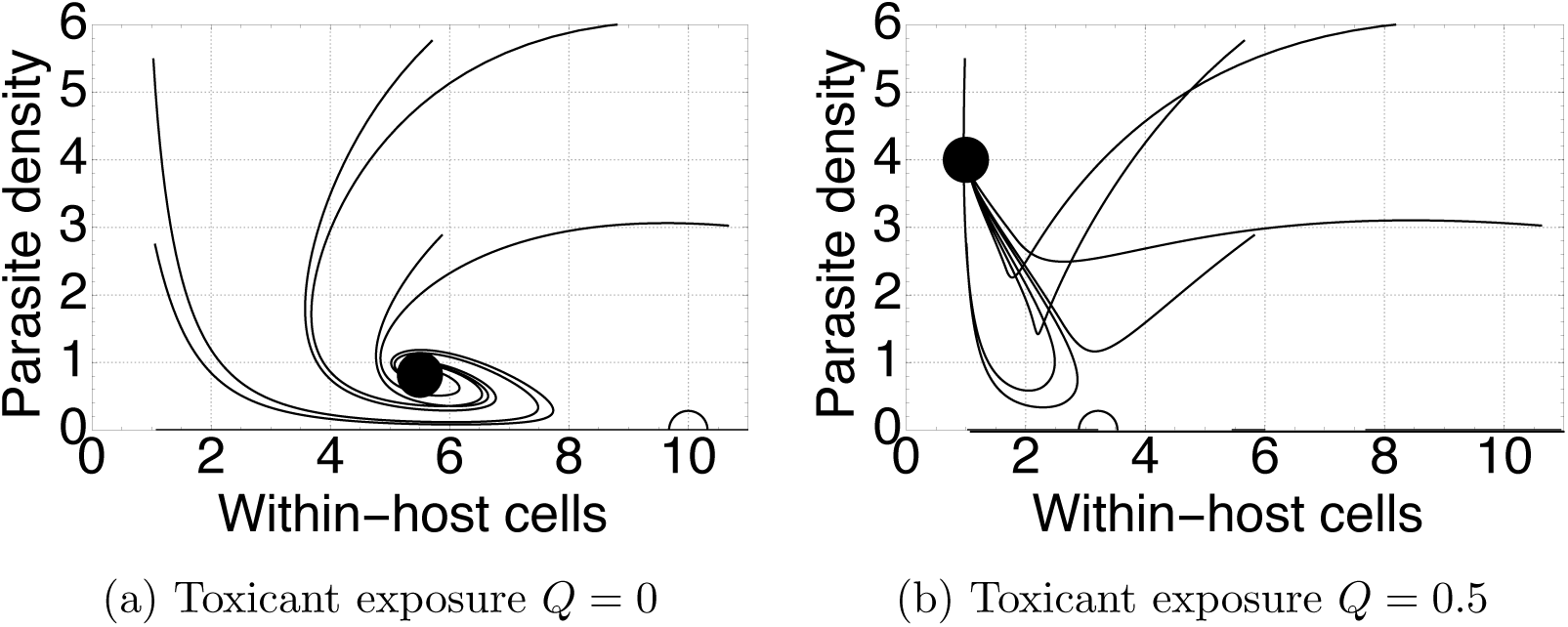
The convergence of the total density of within-host cells and parasites under no toxicant exposure (a) *Q* = 0, and small amounts of toxicant exposure (b) *Q* = 0.5. Black dots show the stable endemic equilibrium, white dots show the unstable disease-free equilibria and lines show the convergence from initial conditions. Parameters as in Table 2 and we assume an initial immune response (*Z* = 10) and an initial amount of within-host cells (*X >* 0), and either zero or positive parasite density (*Y* ≥ 0).

### 3.3. Absence of toxicant lethality

In this case, we consider the absence of a direct lethal toxicant effect, therefore assuming that toxicant exposure only impairs the immune system and does not cause direct mortality. This changes the mechanism by which organisms become infected under increasing toxicant exposure. As before the immune system is inhibited leaving the organism vulnerable to attack by parasites. However after reaching a maximum infected threshold, the health status of the individual remains constant regardless of the amount of toxicant exposure (Figure 4.a). The individual remains highly infected (Figure 4.b) and an increasing exposure to the toxicant no longer causes further damage to organism health status.

**Figure 4:**
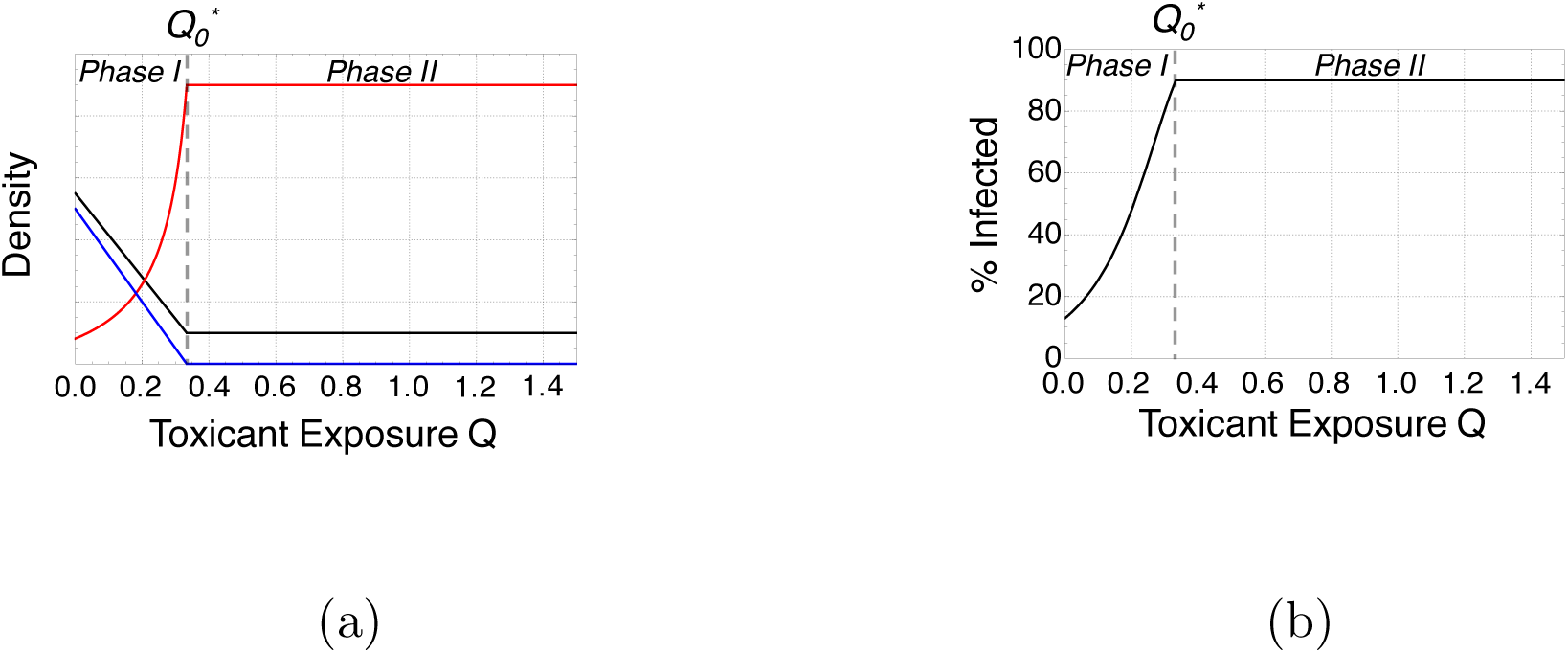
The mechanism of infection under increasing toxicant exposure, under only the immunosuppression of the toxicant effect. In (a), the total density of immune function (blue), parasite load (red) and within-host cells (black) change as an individual is subject to higher toxicant loads. In (b), the total % parasite infection (black) changes as the toxicant load is increased. Parameters as in Table 2.

### 3.4. Aggressive direct mortality

It is worth noting that condition (3) is necessary to explore the interaction between toxicant immunosuppression and the immune system. If this were not the case, for example if the parameter *r* becomes large we would see a situation where the toxicant acts too aggressively upon the host and causes the parasite infection to be killed off (similar to phase II under the original assumption) and following this the within-host cells are destroyed. The immune system remains intact as the direct effect of the toxicant on production of within-host cells is greater than the immune effect. We again see three distinct phases as we increase the toxicant from low levels to high (Figure 5a). However now the toxicant exposure is more prominent and reduces both parasite and within-host cells, stopping the infection from spreading quickly (Figure 5b). In this situation we also see a somewhat contradictory phase 3 in which the host has neither parasite or within-host cells but a small amount of immunity. This result demonstrates the necessity of our original condition.

**Figure 5:**
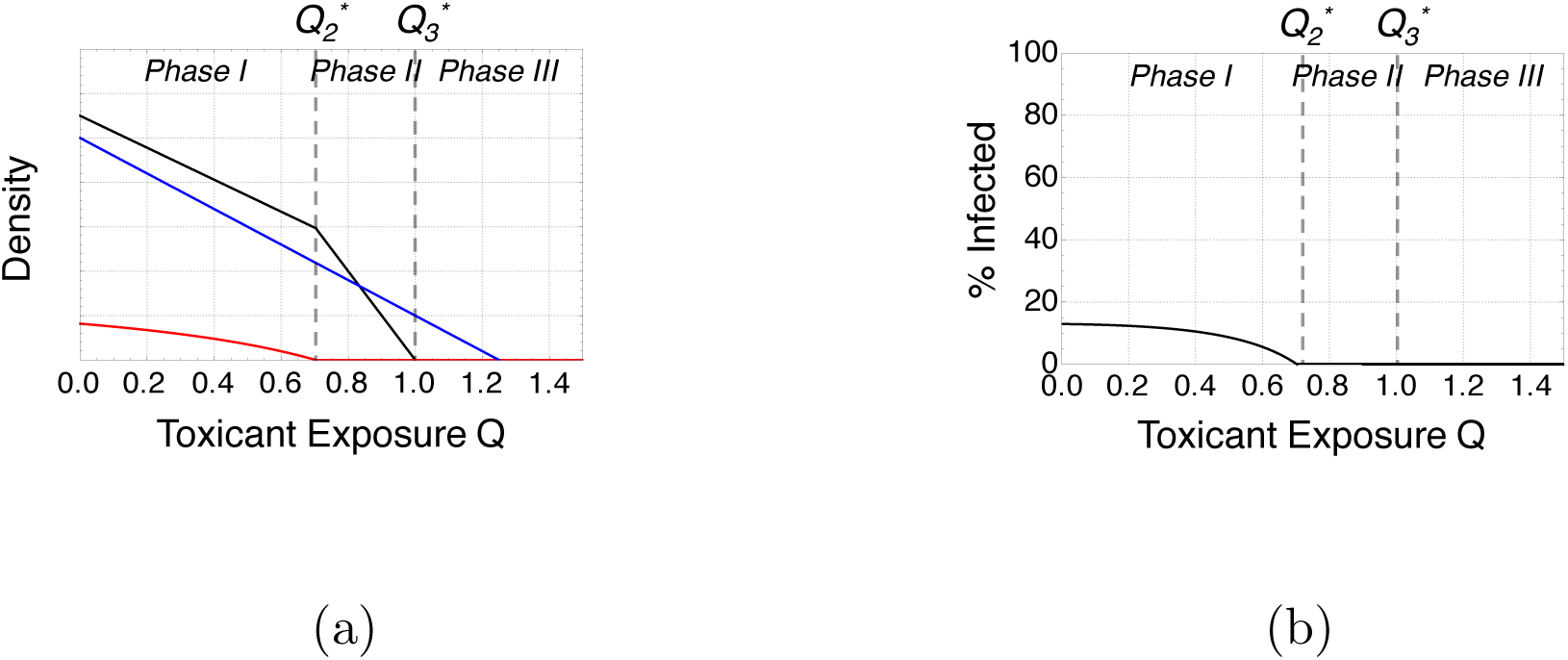
The mechanism of parasite infection under increasing toxicant exposure with aggressive direct mortality. In (a), the total density of immune function (blue), parasite load (red) and within-host cells (black) change as an individual honey bee is subject to higher toxicant loads. In (b), the total % parasite infection (black) changes as the toxicant load is increased. Parameters taken from Table 2, but with a reduced indirect effect *h* = 0.08.

## 4. Discussion

We have shown that interactions between general anthropogenic stress in the form of an immunotoxicant and a parasite can promote within-host infection and reduce health status. The immune response of the host can be divided into three phases of increasing toxicant load; phase I, II and III (Figure 2). In the first phase, sub-lethal doses of the toxicant damage the immune system. This results in suppression of the immune system and hence the individual organism becomes highly infected. In the second phase, intermediate exposure to the toxicant reduces the total density of parasites. In the third phase, the extremely high exposure to the toxicant leads to the loss of within-host cells and eventual mortality of the host.

Through disentangling the individual effects of both lethal and sub-lethal toxicant exposure, we were able to establish the role of each within the breakdown of within-host dynamics. Indirect (sub-lethal) suppression of the immune system causes rapid proliferation of parasites within the host (Figure 3), while direct (lethal) mortality cause both parasites and within-host cells to die. However without the direct effect of the toxicant on the production of new cells, the host remains highly infective (Figure 4). We also predict that an extremely small toxicant exposure can cause the proliferation of a previously manageable infection.

The findings we present in this study shed new light on the poorly understood mechanism by which toxicants seem to interact synergistically with infection to increase mortality risk [65]. In the context of the recent losses to honey bees populations [6, 45, 46], the synergistic immunotoxicant-infection interaction studied here is one example of the recent hypothesis that widespread honey bee losses may be multi-factorial [53, 54, 55, 56]. Synergistic pesticide-infection interactions have been shown to increase mortality risk within honey bees [34, 35]; for example, *Nosema ceranae* infections and thiacloprid, a neonicotinoid pesticide act synergistically to increase individual mortality [37]. The findings we present in this paper propose one explanation of how synergy between these toxicants and infection occur at the within-host level. We show that these sub-lethal effects of anthropogenic stress are potentially more damaging to individual health, aggravating parasitic stress. This is in direct agreement to the positive correlation between low level (field condition) neonicotinoid treatment and increases in parasite and viral in-festations in bees [72, 73]. Infections within individual honey bees can be significantly increased by different levels of low or high sub-lethal pesticides [36]. Indeed, honey bees with undetectable levels of neonicotinoid imidacloprid which are reared in sub-lethal conditions still have increased infection levels [36]. This suggests that even extremely small sub-lethal exposure to pesticide can result in outbreaks of infection. We show that increasing the pesticide exposure by a small amount (*Q >* 0) can result in a transition from a manageable parasite density level to a highly infected individual.

Our results rely upon condition (3) which ensures that the immune response is destroyed before the within-host cells. This condition is crucial to ensuring reasonable behaviour of the model, and it should be noted that the reverse assumption predicts the presence of immunity even after both infected and within-host cells are dead (Figure 5a). We highlight this limitation of our theoretical work but argue that condition (3) is valid since the direct lethality of toxicants only occur at high doses [18] and various immuno-suppressive effects occur from toxicants [39], thus suggesting that toxicants have a greater impact on suppressing the immune system.

The framework provided in this study focuses on the failure of the immune system of an individual organism. However individuals interact within populations causing infection to spread to other susceptible individuals, and these populations have associated interdependent immune defences at both the within-host and between-host level. For example, social immunity involves many behavioural and population-level mechanisms such as social fever, a mechanism by which individuals increase the temperature of the surrounding environment in order to kill parasites [74], guarding, where patrolling guards prevent infected individuals from interacting with healthy individuals [75], hygienic cleaning behavioural traits, by which the population remove diseased or dead individuals [76] and storing antimicrobial food [77]. Hence the main limitation of our framework is that we may have only considered one half of both interdependent within and between-host immunities. Coupling population immunity models in the context of an epidemic alongside our individual immunity framework could further explain the synergistic interactions between toxicants and infection at both the individual and population level. Further theoretical work incorporating these multi-level dynamics could address the gap in understanding honey bee sudden collapse as synergistic stressors in similar ways to other models of colony collapse disorder [78, 79, 80].

This work highlights the need for further studies which focus on synergistic interactions between various stressors at the within-host level. Our theoretical study presents a starting position to think about these synergistic interactions at the within-host level in the context of the immune system of an individual organism. While our model has an inherently simple structure, the addition of the toxicant function can lead to complicated dynamics that are consistent with empirical observations. This framework can stimulate further empirical and theoretical studies which focus on the interaction between toxicant exposure, infection and the immune system at both the social group and individual level.

## 5. Acknowledgments

This work was supported by a Japan Society for the Promotion of Science (JSPS) BRIDGE Fellowship and a University of Sheffield PhD scholarship to R.D.B.

## 7. Authors’ Contributions

All authors conceived the idea for the study, constructed the model and analysed and interpreted the material. R.D.B. wrote the manuscript, with contributions from all authors.

## References

[1]. Nico M. Van Straalen. Ecotoxicology becomes stress ecology. Environmental science & technology, 37:324A–330A, 2003. ISSN 0013-936X. doi:10.1021/es0325720.

[2]. Martin Holmstrup, Anne Mette Bindesbø l, Gertie Janneke Oostingh, Albert Duschl, Volker Scheil, Heinz R. Köhler, Susana Loureiro, Amadeu M.V.M. Soares, Abel L.G. Ferreira, Cornelia Kienle, Almut Gerhardt, Ryszard Laskowski, Paulina E. Kramarz, Mark Bayley, Claus Svendsen, and David J. Spurgeon. Interactions between effects of environmental chemicals and natural stressors: A review, 2010. ISSN 00489697.

[3]. Mieke Jansen, Robby Stoks, Anja Coors, Wendy van Doorslaer, and Luc de Meester. Collateral damage: Rapid exposure-induced evolution of pesticide resistance leads to increased susceptibility to parasites. Evolution, 65:2681–2691, 2011. ISSN 00143820. doi:10.1111/j.1558-5646. 2011.01331.x.

[4]. Alfred Elbert, Matthias Haas, Bernd Springer, Wolfgang Thielert, and Ralf Nauen. Applied aspects of neonicotinoid uses in crop protection. Pest Management Science, 64(11):1099–1105, 2008. ISSN 1526498X. doi:10.1002/ps.1616.

[5]. L. W. Pisa, V. Amaral-Rogers, L. P. Belzunces, J. M. Bonmatin, C. A. Downs, D. Goulson, D. P. Kreutzweiser, C. Krupke, M. Liess, M. McField, C. A. Morrissey, D. A. Noome, J. Settele, N. Simon-Delso, J. D. Stark, J. P. Van der Sluijs, H. Van Dyck, and M. Wiemers. Effects of neonicotinoids and fipronil on non-target invertebrates. Environmental science and pollution research international, 22(1):68–102, 2015. ISSN 16147499. doi:10.1007/s11356-014-3471-x.

[6]. Dave Goulson, Elizabeth Nicholls, Cristina Botias, and Ellen L Rotheray. Bee declines driven by combined stress from parasites, pesticides, and lack of flowers. Science, 347(6229):1255957, 2015. ISSN 1095-9203. doi:10.1126/science.1255957.

[7]. E. C. Oerke and H. W. Dehne. Safeguarding production - Losses in major crops and the role of crop protection. Crop Protection, 23(4): 275–285, 2004. ISSN 02612194. doi:10.1016/j.cropro.2003.10.001.

[8]. European Food Safety Authority. Conclusion on the peer review of the pesticide risk assessment for bees for Clothianidin. EFSA Journal, 11: 1–58, 2013. ISSN 1831-4732. doi:10.2903/j.efsa.2013.3066.

[9]. European Food Safety Authority. Conclusion on the peer review of the pesticide risk assessment for bees for the active substance imidacloprid. EFSA Journal, 11:1–55, 2013. ISSN 1831-4732. doi:10.2903/j.efsa.2013. 3068.

[10]. Michael Gross. EU ban puts spotlight on complex effects of neonicotinoids. Current Biology, 23(11):R462–R464, 2013. ISSN 09609822. doi:10.1016/j.cub.2013.05.030.

[11]. C. H. Krupke, J. D. Holland, E. Y. Long, and B. D. Eitzer. Planting of neonicotinoid-treated maize poses risks for honey bees and other non-target organisms over a wide area without consistent crop yield benefit. Journal of Applied Ecology, 2017. ISSN 13652664. doi:10.1111/1365-2664.12924.

[12]. H. C. J. Godfray, T. Blacquiere, L. M. Field, R. S. Hails, G. Petrokofsky, S. G. Potts, N. E. Raine, a. J. Vanbergen, and a. R. McLean. A restatement of the natural science evidence base concerning neonicotinoid insecticides and insect pollinators. Proceedings of the Royal Society B: Biological Sciences, 281:20140558, 2014. ISSN 1471-2954. doi:10.1098/rspb.2014.0558. URL http://www.ncbi.nlm.nih.gov/pubmed/24850927.

[13]. H. Charles J. Godfray, Tjeerd Blacquiere, Linda M. Field, Rosemary S. Hails, Simon G. Potts, Nigel E. Raine, Adam J. Vanbergen, An-gela R. McLean, H. Charles J. Godfray, Tjeerd Blacquie, Simon G. Potts, Nigel E. Raine, Adam J. Vanbergen, and Angela R. McLean. A restatement of recent advances in the natural science evidence base concerning neonicotinoid insecticides and insect pollinators. Proceedings of the Royal Society B: Biological Sciences, 282:20151821, 2015. ISSN 0962-8452. doi:10.1098/rspb.2015.1821. URL http://rspb.royalsocietypublishing.org/content/282/1818/20151821.

[14]. J L Bernal, E Garrido-Bailon, M J Del Nozal, a V Gonzalez-Porto, R Martin-Hernandez, J C Diego, J J Jimenez, J L Bernal, and M Higes. Overview of pesticide residues in stored pollen and their potential effect on bee colony (Apis mellifera) losses in Spain. Journal of economic entomology, 103(6):1964–1971, 2010. ISSN 0022-0493. doi:10.1603/ EC10235.

[15]. Dave Goulson. An overview of the environmental risks posed by neonicotinoid insecticides. Journal of Applied Ecology, 50(4):977–987, 2013. ISSN 00218901. doi:10.1111/1365-2664.12111.

[16]. Cristina Botias, Arthur David, Julia Horwood, Alaa Abdul-Sada, Elizabeth Nicholls, Elizabeth Hill, and Dave Goulson. Neonicotinoid Residues in Wildflowers, a Potential Route of Chronic Exposure for Bees. Environmental Science and Technology, 49:12731–12740, 2015. ISSN 15205851. doi:10.1021/acs.est.5b03459.

[17]. T Blacquiere, G. Smagghe, C. A. Van Gestel, and V. Mommaerts. Neonicotinoids in bees: a review on concentrations, side-effects and risk assessment. Ecotoxicology, 21(4):973–992, 2012.

[18]. T. Iwasa, N. Motoyama, J. T. Ambrose, and R. M. Roe. Mechanism for the differential toxicity of neonicotinoid insecticides in the honey bee, Apis mellifera. Crop Protection, 23(5):371–378, 2004.

[19]. Hongsheng Pan, Yongqiang Liu, Bing Liu, Yanhui Lu, Xiaoyong Xu, Xuhong Qian, Kongming Wu, and Nicolas Desneux. Lethal and sublethal effects of cycloxaprid, a novel cis-nitromethylene neonicotinoid insecticide, on the mirid bug Apolygus lucorum. Journal of Pest Science, 87:731–738, 2014. ISSN 16124758. doi:10.1007/s10340-014-0610-6.

[20]. Ran Wang, Wei Zhang, Wunan Che, Cheng Qu, Fengqi Li, Nicolas Desneux, and Chen Luo. Lethal and sublethal effects of cyantraniliprole, a new anthranilic diamide insecticide, on Bemisia tabaci (Hemiptera: Aleyrodidae) MED. Crop Protection, 91:108–113, 2017. ISSN 02612194. doi:10.1016/j.cropro.2016.10.001.

[21]. Richard J Gill, Oscar Ramos-Rodriguez, and Nigel E Raine. Combined pesticide exposure severely affects individual- and colony-level traits in bees. Nature, 491(7422):105–8, November 2012. ISSN 1476-4687. doi:10.1038/nature11585. URL http://www.pubmedcentral.nih.gov/articlerender.fcgi?artid=3495159&tool=pmcentrez&rendertype=abstract.

[22]. Richard J. Gill and Nigel E. Raine. Chronic impairment of bumblebee natural foraging behaviour induced by sublethal pesticide exposure. Functional Ecology, 28:1459–1471, 2014. ISSN 13652435. doi:10.1111/1365-2435.12292.

[23]. Hannah Feltham, Kirsty Park, and Dave Goulson. Field realistic doses of pesticide imidacloprid reduce bumblebee pollen foraging efficiency. Ecotoxicology, 23:317–323, 2014. ISSN 15733017. doi:10.1007/s10646-014-1189-7.

[24]. Dara A. Stanley, Avery L. Russell, Sarah J. Morrison, Catherine Rogers, and Nigel E. Raine. Investigating the impacts of field-realistic exposure to a neonicotinoid pesticide on bumblebee foraging, homing ability and colony growth. Journal of Applied Ecology, 53:1440–1449, 2016. ISSN 13652664. doi:10.1111/1365-2664.12689.

[25]. Peng Han, Chang Ying Niu, Chao Liang Lei, Jin Jie Cui, and Nicolas Desneux. Quantification of toxins in a Cry1Ac + CpTI cotton cultivar and its potential effects on the honey bee Apis mellifera L. Ecotoxicology, 19(8):1452–1459, 2010. ISSN 09639292. doi:10.1007/s10646-010-0530-z.

[26]. Axel Decourtye, Eric Lacassie, and Minh Ha Pham-Delegue. Learning performances of honeybees (Apis mellifera L) are differentially affected by imidacloprid according to the season. Pest Management Science, 59 (3):269–278, 2003. ISSN 1526498X. doi:10.1002/ps.631.

[27]. SM Williamson and GA Wright. Exposure to multiple cholinergic pesticides impairs olfactory learning and memory in honeybees. The Journal of experimental biology, 216(10):1799–1807, 2013.

[28]. Axel Decourtye, Catherine Armengaud, Michel Renou, James Devillers, Sophie Cluzeau, Monique Gauthier, and Minh Ha Pham-Delegue. Imidacloprid impairs memory and brain metabolism in the honeybee (Apis mellifera L.). Pesticide Biochemistry and Physiology, 78(2):83–92, 2004. ISSN 00483575. doi:10.1016/j.pestbp.2003.10.001.

[29]. Penelope R. Whitehorn, Stephanie O'connor, Felix L. Wackers, and Dave Goulson. Neonicotinoid Pesticide Reduces Bumble Bee Colony Growth and Queen Production. Science, 336:351–352, 2012. ISSN 1526498X. doi:10.1002/ps.1616.

[30]. Maj Rundlöf, Georg K. S. Andersson, Riccardo Bommarco, Ingemar Fries, Veronica Hederström, Lina Herbertsson, Ove Jonsson, Björn K. Klatt, Thorsten R. Pedersen, Johanna Yourstone, and Henrik G. Smith. Seed coating with a neonicotinoid insecticide negatively affects wild bees. Nature, 521:77–80, 2015. ISSN 0028-0836. doi:10.1038/nature14420. URL http://www.nature.com/doifinder/10.1038/nature14420.

[31]. Geoffrey R. Williams, Aline Troxler, Gina Retschnig, Kaspar Roth, Orlando Yañez, Dave Shutler, Peter Neumann, and Laurent Gauthier. Neonicotinoid pesticides severely affect honey bee queens. Scientific Reports, 5:14621, 2015. ISSN 2045-2322. doi:10.1038/srep14621. URL http://www.pubmedcentral.nih.gov/articlerender.fcgi?artid=4602226&tool=pmcentrez&rendertype=abstract.

[32]. En Cheng Yang, Hui Chun Chang, Wen Yen Wu, and Yu Wen Chen. Impaired Olfactory Associative Behavior of Honeybee Workers Due to Contamination of Imidacloprid in the Larval Stage. PLoS ONE, 7(11): e49472, 2012. ISSN 19326203. doi:10.1371/journal.pone.0049472.

[33]. Ken Tan, Weiwen Chen, Shihao Dong, Xiwen Liu, Yuchong Wang, and James C Nieh. A neonicotinoid impairs olfactory learning in Asian honey bees (Apis cerana) exposed as larvae or as adults. Scientific reports, 5: 10989, 2015. ISSN 2045-2322. doi:10.1038/srep10989.

[34]. Cedric Alaux, fr Jean Luc Brunet, Claudia Dussaubat, Fanny Mondet, Sylvie Tchamitchan, Marianne Cousin, Julien Brillard, Aurelie Baldy, Luc P. Belzunces, and Yves Le Conte. Interactions between Nosema microspores and a neonicotinoid weaken honeybees (Apis mellifera). Environmental Microbiology, 12(3):774–782, 2010. ISSN 14622912. doi:10.1111/j.1462-2920.2009.02123.x.

[35]. Cyril Vidau, Marie Diogon, Julie Aufauvre, Régis Fontbonne, Bernard Vigues, Jean Luc Brunet, Catherine Texier, David G. Biron, Nicolas Blot, Hicham Alaoui, Luc P. Belzunces, and Frédéric Delbac. Exposure to sublethal doses of fipronil and thiacloprid highly increases mortality of honeybees previously infected by nosema ceranae. PLoS ONE, 6(6): e21550, 2011. ISSN 19326203. doi:10.1371/journal.pone.0021550.

[36]. JS Pettis, J Johnson, and G Dively. Pesticide exposure in honey bees results in increased levels of the gut pathogen Nosema. Naturwissenschaften, 99(2):153–158, 2012.

[37]. Vincent Doublet, Maureen Labarussias, Joachim R. de Miranda, Robin F. A. Moritz, and Robert J. Paxton. Bees under stress: sublethal doses of a neonicotinoid pesticide and pathogens interact to elevate honey bee mortality across the life cycle. Environmental Microbiology, 17(4): 969–983, April 2015. ISSN 14622912. doi:10.1111/1462-2920.12426.

[38]. Anja Coors, Ellen Decaestecker, Mieke Jansen, and Luc De Meester. Pesticide exposure strongly enhances parasite virulence in an invertebrate host model. Oikos, 117:1840–1846, 2008. ISSN 00301299. doi:10.1111/j.1600-0706.2008.17028.x.

[39]. R. R. James and J. Xu. Mechanisms by which pesticides affect insect immunity. Journal of Invertebrate Pathology, 109(2):175–182, 2012. ISSN 00222011. doi:10.1016/j.jip.2011.12.005.

[40]. Patricia de Azambuja, E. S. Garcia, N. A. Ratcliffe, and J. David Warthen. Immune-depression in Rhodnius prolixus induced by the growth inhibitor, azadirachtin. Journal of Insect Physiology, 37(10): 771–777, 1991. ISSN 00221910. doi:10.1016/0022-1910(91)90112-D.

[41]. Annely Brandt, Anna Gorenflo, Reinhold Siede, Marina Meixner, and Ralph Buchler. The neonicotinoids thiacloprid, imidacloprid, and clothianidin affect the immunocompetence of honey bees (Apis mellifera L.). Journal of Insect Physiology, 86:40–47, 2016. ISSN 00221910. doi:10.1016/j.jinsphys.2016.01.001.

[42]. A. Zibaee and A. R. Bandani. Effects of Artemisia annua L.(Asteracea) on the digestive enzymatic profiles and the cellular immune reactions of the Sunn pest, Eurygaster integriceps (Heteroptera: Scutellaridae), against Beauveria bassiana. Bulletin of entomological research, 100(02): 185–196, 2010.

[43]. J-M Delpuech, F Frey, and Y Carton. Action of insecticides on the cellular immune reaction of drosophila melanogaster against the parasitoid leptopilina boulardi. Environmental Toxicology & Chemistry, 15(12): 2267–2271, 1996. ISSN 07307268.

[44]. Gennaro Di Prisco, Valeria Cavaliere, Desiderato Annoscia, Paola Varricchio, Emilio Caprio, Francesco Nazzi, Giuseppe Gargiulo, and Francesco Pennacchio. Neonicotinoid clothianidin adversely affects insect immunity and promotes replication of a viral pathogen in honey bees. Proceedings of the National Academy of Sciences, 110(46):18466– 18471, 2013. ISSN 1091-6490. doi:10.1073/pnas.1314923110.

[45]. Romée van der Zee, Lennard Pisa, Sreten Andonov, Robert Brodschnei-der, Jean-Daniel Charrière, Robert Chlebo, Mary F Coffey, Karl Crailsheim, Bjø rn Dahle, Anna Gajda, Alison Gray, Marica M Drazic, Mariano Higes, Lassi Kauko, Aykut Kence, Meral Kence, Nicola Kezic, Hrisula Kiprijanovska, Jasna Kralj, Preben Kristiansen, Raquel Martin Hernandez, Franco Mutinelli, Bach Kim Nguyen, Christoph Otten, Asli özkirim, Stephen F Pernal, Magnus Peterson, Gavin Ramsay, Violeta Santrac, Victoria Soroker, Graz?yna Topolska, Aleksandar Uzunov, Flemming Vejsnæs, Shi Wei, and Selwyn Wilkins. Managed honey bee colony losses in Canada, China, Europe, Israel and Turkey, for the winters of 2008-9 and 2009-10. Journal of Apicultural Research, 51(1): 100–114, 2012. ISSN 20786913. doi:10.3896/IBRA.1.51.1.12.

[46]. Dennis VanEngelsdorp, Dewey Caron, Jerry Hayes, Robyn Underwood, Mark Henson, Karen Rennich, Angela Spleen, Michael Andree, Robert Snyder, Kathleen Lee, Karen Roccasecca, Michael Wilson, James Wilkes, Eugene Lengerich, and Jeffery Pettis. A national survey of managed honey bee 2010–11 winter colony losses in the USA: results from the Bee Informed Partnership. Journal of Apicultural Research, 51(1): 115–124, 2012. ISSN 0021-8839. doi:10.3896/IBRA.1.51.1.14.

[47]. Sven Lautenbach, Ralf Seppelt, Juliane Liebscher, and Carsten F. Dormann. Spatial and temporal trends of global pollination benefit. PLoS ONE, 7:e35954, 2012. ISSN 19326203. doi:10.1371/journal.pone. 0035954.

[48]. N Gallai, JM Salles, J Settele, and BE Vaissiere. Economic valuation of the vulnerability of world agriculture confronted with pollinator decline. Ecological economics, 68(3):810–821, 2009.

[49]. Mike H. Allsopp, Willem J. de Lange, and Ruan Veldtman. Valuing insect pollination services with cost of replacement. PLoS ONE, 3(9): e3128, 2008. ISSN 19326203. doi:10.1371/journal.pone.0003128.

[50]. R A Morse and N W Calderone. The value of honey bees as pollinators of US crops in 2000. Bee Culture, 128(3):1–15, 2000.

[51]. AM Klein, BE Vaissiere, JH Cane, I Steffan-Dewenter, SA Cunningham, C Kremen, and T Tscharntke. Importance of pollinators in changing landscapes for world crops. Proceedings of the Royal Society of London B: Biological Sciences, 274(1608):303–313, 2007. ISSN 0962-8452. doi:10.1098/rspb.2006.3721.

[52]. Jordi Bascompte, Pedro Jordano, and Jens M Olesen. Asymetric Coevolutionary Networks Facilitate Biodiversity Maintenance. Science, 312 (5772):431–433, 2006. ISSN 1095-9203. doi:10.1126/science.1123412.

[53]. SG Potts, JC Biesmeijer, and C Kremen. Global pollinator declines: trends, impacts and drivers. Trends in ecology & evolution, 25(6):345–353, 2010.

[54]. A J Vanbergen. Threats to an ecosystem service: pressures on pollinators. Frontiers in Ecology and the Environment, 11(5):251–259, 2013.

[55]. P Neumann and N L Carreck. Honey bee colony losses. Journal of Apicultural Research, 49:1–6, 2010. ISSN 20786913. doi:10.3896/IBRA. 1.49.1.01.

[56]. FLW Ratnieks and NL Carreck. Clarity on honey bee collapse? Science, 327(5962):152–3, January 2010. ISSN 1095-9203. doi:10.1126/science. 1185563.

[57]. Ola Lundin, Maj Rundlöf, Henrik G. Smith, Ingemar Fries, and Riccardo Bommarco. Neonicotinoid insecticides and their impacts on bees: A systematic review of research approaches and identification of knowledge gaps. PLoS ONE, 10, 2015. ISSN 19326203. doi:10.1371/journal.pone. 0136928.

[58]. Peter Rosenkranz, Pia Aumeier, and Bettina Ziegelmann. Biology and control of Varroa destructor. Journal of Invertebrate Pathology, 103: S96–S119, 2010. ISSN 0022-2011. doi:10.1016/j.jip.2009.07.016.

[59]. D Sammataro, U Gerson, and G Needham. Parasitic mites of honey bees: life history, implications, and impact. Annual review of entomology, 45 (1):519–548, 2000. ISSN 0066-4170. doi:10.1146/annurev.ento.45.1.519.

[60]. A C. Highfield, A El Nagar, LCM Mackinder, LMLJ Nöel, MJ. Hall, SJ. Martin, and DC. Schroeder. Deformed wing virus implicated in overwintering honeybee colony losses. Applied and Environmental Microbiology, 75(22):7212–7220, 2009. ISSN 00992240. doi:10.1128/AEM.02227-09.

[61]. Eugene V. Ryabov, Graham R. Wood, Jessica M. Fannon, Jonathan D. Moore, James C. Bull, Dave Chandler, Andrew Mead, Nigel Burroughs, and David J. Evans. A Virulent Strain of Deformed Wing Virus (DWV) of Honeybees (Apis mellifera) Prevails after Varroa destructor-Mediated, or In Vitro, Transmission. PLoS Pathogens, 10, 2014. ISSN 15537374. doi:10.1371/journal.ppat.1004230.

[62]. J Moore, A Jironkin, D Chandler, N Burroughs, DJ Evans, and EV. Ryabov. Recombinants between Deformed wing virus and Varroa destructor virus-1 may prevail in Varroa destructor-infested honeybee colonies. Journal of General Virology, 92(1):156–161, 2011. ISSN 00221317. doi:10.1099/vir.0.025965-0.

[63]. Mariano Higes, Raquel Martín-Hernàndez, Encarna Garrido-Bailon, Amelia V. González-Porto, Pilar García-Palencia, Aranzazu Meana, María J. del Nozal, R. Mayo, and José L. Bernal. Honeybee colony collapse due to Nosema ceranae in professional apiaries. Environmental Microbiology Reports, 1(2):110–113, 2009. ISSN 17582229. doi:10.1111/j.1758-2229.2009.00014.x.

[64]. M Higes and A Meana. Nosema ceranae (Microsporidia), a controversial 21st century honey bee pathogen. Environmental microbiology reports, 5(1):17–29, 2013.

[65]. James E. Cresswell, Nicolas Desneux, and Dennis VanEngelsdorp. Dietary traces of neonicotinoid pesticides as a cause of population declines in honey bees: An evaluation by Hill's epidemiological criteria. Pest Management Science, 68(6):819–827, 2012. ISSN 1526498X. doi:10.1002/ps.3290.

[66]. Martin Nowak and Robert M. May. Virus dynamics: mathematical principles of immunology and virology: mathematical principles of immunology and virology. Oxford University Press, UK, 2000.

[67]. Y. Chen, J. D. Evans, I. B. Smith, and J. S. Pettis. Nosema ceranae is a long-present and wide-spread microsporidian infecton of the European honey bee (Apis mellifera) in the United States. Journal of Invertebrate Pathology, 97(2):186–188, 2008. doi:http://dx.doi.org/10.1016/j.jip.2007.07.010. URL http://www.sciencedirect.com/science/article/pii/S002220110700153X.

[68]. Hoda M. Nasr, Mohamed E I Badawy, and Entsar I. Rabea. Toxicity and biochemical study of two insect growth regulators, buprofezin and pyriproxyfen, on cotton leafworm Spodoptera littoralis. Pesticide Biochemistry and Physiology, 98(2):198–205, 2010. ISSN 00483575. doi:10.1016/j.pestbp.2010.06.007.

[69]. Z Ma, X Han, J Feng, G Li, and X ZHANG. Effects of Terpinen-4-ol on Four Metabolic Enzymes and Polyphenol Oxidase (PPO) in Mythimna separta Walker. Agricultural Sciences in China, 7:726–730, 2008. ISSN 16712927. doi:10.1016/S1671-2927(08)60107-8.

[70]. Ender Büyükgüzel. Evidence of oxidative and antioxidative responses by Galleria mellonella larvae to malathion. Journal of economic entomology, 102(1):152–159, 2009. ISSN 00220493. doi:10.1603/029.102.0122.

[71]. S Alizon and M van Baalen. Acute or Chronic? Within-Host Models with Immune Dynamics, Infection Outcome, and Parasite Evolution. American Naturalist, 172:E244–E256, 2008. ISSN 0003-0147. doi:10. 1086/592404.

[72]. Galen P. Dively, Michael S. Embrey, Alaa Kamel, David J. Hawthorne, and Jeffery S. Pettis. Assessment of chronic sublethal effects of imidacloprid on honey bee colony health. PLoS ONE, 10, 2015. ISSN 19326203. doi:10.1371/journal.pone.0118748.

[73]. Francisco Sánchez-Bayo, Dave Goulson, Francesco Pennacchio, Francesco Nazzi, Koichi Goka, and Nicolas Desneux. Are bee diseases linked to pesticides? - A brief review, 2016. ISSN 18736750.

[74]. Philip T. Starks, Caroline A. Blackie, and P. T. Thomas D Seeley. Fever in honeybee colonies. Naturwissenschaften, 87(5):229–231, 2000. ISSN 00281042. doi:10.1007/s001140050709.

[75]. Keith D. Waddington and Walter C. Rothenbuhler. Behaviour Associated with Hairless-Black Syndrome of Adult Honeybees. Journal of Apicultural Research, 15(1):35–41, 1976. ISSN 0021-8839. doi:10.1080/00218839.1976.11099831.

[76]. Marla Spivak and M. Gilliam. Hygienic behaviour of honey bees and its application for control of brood diseases and Varroa. Part II. Studies on hygienic behaviour since the Rothenbuhler era. Bee World, 79:169–186, 1998. ISSN 0005772X. doi:10.1080/0005772X.1998.11099394.

[77]. R. A. Morse and K. Flottum. Honey bee pests, predators, and diseases. AI Root Company, Medina, OH, 3rd edition, 1997. ISBN 0936028106.

[78]. J Bryden, RJ Gill, RAA Mitton, NE Raine, and VAA Jansen. Chronic sublethal stress causes bee colony failure. Ecology letters, 16(12):1463– 1469, 2013. ISSN 1461-0248. doi:10.1111/ele.12188.

[79]. Ross D. Booton, Yoh Iwasa, James A.R. Marshall, and Dylan Z. Childs. Stress-Mediated Allee Effects Can Cause the Sudden Collapse of Honey Bee Colonies. Journal of Theoretical Biology, 420:213–219, 2017. ISSN 00225193. doi:10.1016/j.jtbi.2017.03.009. URL http://linkinghub.elsevier.com/retrieve/pii/S0022519317301212.

[80]. Clint J Perry, Eirik Søvik, Mary R Myerscough, and Andrew B Barron. Rapid behavioral maturation accelerates failure of stressed honey bee colonies. Proceedings of the National Academy of Sciences of the United States of America, 112:3427–32, 2015. ISSN 1091-6490. doi:10.1073/pnas.1422089112. URL http://www.pnas.org/content/112/11/3427.short.

